# Quantitative Molecular Imaging of Breast Microcalcification Composition using Photon-Counting Spectral Computed Tomography

**DOI:** 10.1101/2020.09.16.300509

**Authors:** Tyler E. Curtis, Ryan K. Roeder

**Affiliations:** Department of Aerospace and Mechanical Engineering, Bioengineering Graduate Program, University of Notre Dame, Notre Dame, IN 46556, USA

**Keywords:** Breast Microcalcifications, Calcium Phosphate, Calcium Oxalate, Hydroxyapatite, Molecular Imaging, Quantitative Material Decomposition, Photon-Counting Spectral Computed Tomography

## Abstract

Mammographic screening for breast cancer is unable to distinguish molecular differences between hydroxyapatite (HA) microcalcifications (μcals) that are associated with malignancy and calcium oxalate (CaOx) μcals that are benign. Therefore, the objective of this study was to investigate quantitative material decomposition of model breast μcals of clinically-relevant composition and size using spectral photon-counting computed tomography (PCCT). Model μcals composed of HA, CaOx, and dicalcium phosphate (DCP) were treated as materials containing spatially coincident elemental compositions of calcium (Ca), phosphorus (P), and oxygen (O). Elemental decomposition was performed using constrained maximum-likelihood estimation in the image domain. Images were acquired with a commercial, preclinical PCCT system (MARS Bioimaging) with five energy bins selected to maximize counts at low photon energies and spectral differences between Ca and P. Elemental concentrations of Ca and P within the each μcal composition were accurately identified and quantified with a root-mean-squared error < 12%. HA and CaOx μcals, < 1 mm is size, were accurately discriminated by the measured P content with an area under the receiver operating characteristic curve (AUC) > 0.9. The mole fraction of P, P/(Ca+P), was able to discriminate all three μcal compositions with AUC > 0.8 for μcals < 1 mm is size and AUC = 1 for μcals > 2 mm in size. The overall accuracy for the classification of μcal types and quantification of P was robust against different assumptions in the elemental decomposition calibration, but quantification of Ca was improved with assumptions that most accurately accounted for the molar volume of each element within μcal compositions. Thus, PCCT enabled quantitative molecular imaging of breast μcal composition, which is not possible with current clinical molecular imaging modalities.

## 1. Introduction

Breast cancer is the most widely diagnosed type of cancer, and the second leading cause of cancer-related deaths, among women in the United States (Siegel *et al*. 2019). Early radiographic detection by mammography is currently the best available means for reducing breast cancer mortality (Smith *et al*. 2019, Van Ourti *et al*. 2020). Microcalcifications (μcals) are the most common abnormality detected by mammography, coincident with 30-50% of all breast lesions and 85-95% of all cases of ductal carcinoma *in situ* (Cheng *et al*. 2003, Gülsün *et al*. 2003, Henrot *et al*. 2014, Millis *et al*. 1976). Two types of μcals commonly present in breast tissue (Bonfiglio *et al*. 2018, Frappart *et al*. 1984, Frappart *et al*. 1987, Haka *et al*. 2002). Type I μcals are composed of calcium oxalate (CaOx), CaC_2_O_4_, and are benign. Type II μcals are composed of hydroxyapatite (HA), Ca_5_(PO_4_)_3_OH, and are associated with malignant lesions. Both types of μcals are detected radiographically by high X-ray contrast relative to adjacent soft tissue. Therefore, radiologists currently use qualitative features – such as the μcal morphology, size, anatomic location and clustering – to distinguish suspicious Type II μcals from benign Type I μcals (Bonfiglio *et al*. 2018). After the observation of a suspicious μcal in a mammogram, a biopsy is required for definitive diagnosis, but malignancy is only confirmed in only 20-30% of biopsies (Kim *et al*. 2020, White *et al*. 2001). Thus, mammographic screening results in false-positives, unnecessary biopsies, and overdiagnosis (Løberg *et al*. 2015, Oeffinger *et al*. 2015, Siu 2016).

Spectral or multi-energy X-ray imaging methods – such as dual-energy computed tomography (DECT) or mammography and photon-counting CT (PCCT) – have recently enabled quantitative imaging of material composition by leveraging characteristic differences in X-ray attenuation across the incident photon energy spectrum (Jacobsen *et al*. 2020, McCollough *et al*. 2020, Willemink *et al*. 2018). Conventional CT and mammography utilize an energy integrating detector (EID) which sums all photon interactions across the observable energy spectrum into a single image, ignoring spectral and thus material-specific information. DECT partially leverages spectral information by exposing an EID to two different but overlapping photon energy spectra, such that differences in X-ray attenuation can be used to identify (Johnson *et al*. 2007) and quantify (Yu *et al*. 2018) material compositions, but material identification in DECT is typically limited to two material components. Assumptions can be made in the material decomposition to allow up to three material components, but can also lead to increased error. Photon-counting detectors (PCDs) enable spectral information to be acquired from a single source spectra in multiple, distinct energy bins by thresholding direct current peaks associated with the energy of photon interactions (Willemink *et al*. 2018). Therefore, PCCT can identify (Anderson *et al*. 2010, Cormode *et al*. 2010) and quantify (Alessio and MacDonald 2013, Cormode *et al*. 2017, McGinnity *et al*. 2016, Perrson *et al*. 2014, Schlomka *et al*. 2008) multiple discrete and even mixed (Curtis and Roeder 2019, Symons *et al*. 2017) material compositions simultaneously.

Several recent studies have investigated the feasibility of discriminating calcium compositions associated with breast μcals and renal stones using spectral X-ray imaging methods. HA and CaOx renal stones, >3 mm in size, were identified at an area under the receiver operating characteristic curve (AUC) of ∼0.8 using DECT with either an EID or PCD (Ferrero *et al*. 2018). The feasibility of discriminating model HA and CaOx breast μcals was demonstrated by numerical and analytical simulations of dual energy mammography with a PCD (Ghammraoui *et al*. 2017, 2019, Kim *et al*. 2019, Martini *et al*. 2017). Experimental validations reported AUC ranging 0.76-0.88 (Ghammraoui *et al*. 2019), or measured differences in an “effective” calcium-to-phosphorus ratio (Ca/P) with a coefficient of variation ranging ∼5-20% (Martini *et al*. 2017), for variations in the phantom and μcal thickness. Importantly, the preceding dual energy methods all utilized relative differences in the measured signal or contrast, normalized to noise or background, between a low and high energy bin or acquisition, and were therefore unable to determine the μcal elemental composition. The discrimination of HA and CaOx by multi-energy PCCT was investigated by comparing the measured contrast-to-noise ratio across several photon energy bins, but material decomposition was not performed (Kirkbride *et al*. 2017).

Therefore, the objective of this study was to investigate quantitative material decomposition of model breast μcals of clinically-relevant composition and size using PCCT. Model μcals composed of HA (type II, associated with malignancy), CaOx (type I, benign), and dicalcium phosphate (DCP, control) were treated as materials containing mixed elemental compositions of calcium (Ca), phosphorus (P), and oxygen (O). Elemental concentration maps, and the mole fraction of P, P/(Ca+P), was used to identify and discriminate between each type of μcal.

## 2. Materials and Methods

### 2.1. Homogenous Liquid Phantom

Elemental decomposition of Ca and P was first investigated with a homogeneous liquid phantom containing mixtures of calcium chloride and phosphoric acid in varying molar ratios.

#### 2.1.1. Calibration and Sample Phantoms

Calibration phantoms comprising calcium chloride dihydrate, CaCl_2_·2H_2_O (Sigma Aldrich, ≥ 99.9%), and phosphoric acid, H_3_PO_4_ (Sigma Aldrich, > 85 wt%) solutions were prepared with 8 concentrations each, ranging 0 to 4.8 M (192 g/L) Ca and 0 to 4 M (124 g/L) P, respectively, in deionized (DI) water containing 1% agarose (Thermo Scientific, molecular biology grade) (Table 1). Calibration concentrations were chosen to significantly exceed estimated sample concentrations to achieve greater accuracy and less variability in image-based material decomposition by maximizing the signal-to-noise ratio for each component (Curtis and Roeder, 2017). A sample phantom comprising mixtures of calcium chloride and phosphoric acid was prepared at 8 concentrations ranging from 0 to 2.44 M (97.8 g/L) Ca and 2.17 M (67.2 g/L) to 0 M P, respectively, in DI water containing 1% agarose (Table 1). The calibration phantom was imaged once and the sample phantom was imaged in ten separate acquisitions, to create ten imaging replicates of each mixed composition of Ca and P. For both the calibration and sample phantoms, Eppendorf tubes (5 mm inner diameter) containing each discrete or mixed composition and concentration were placed in a modular acrylic phantom (35 mm outer diameter) for imaging. A polytetrafluoroethylene screw was present at the center of each phantom as part of a clamping mechanism for holding Eppendorf tubes thin the phantom and was excluded from material decomposition.

**Table 1.** Discrete and mixed concentrations of Ca and P within the homogenous liquid calibration and sample phantoms, respectively.

| Phantom | Component | Concentrations (M) |  |  |  |  |  |  |  |
| --- | --- | --- | --- | --- | --- | --- | --- | --- | --- |
| Calibration<br>(discrete Ca and P) | Ca | 0 | 0.69 | 1.37 | 2.05 | 2.74 | 3.43 | 4.11 | 4.80 |
|  | P | 0 | 0.57 | 1.14 | 1.71 | 2.29 | 2.86 | 3.42 | 4.00 |
| Sample<br>(mixed Ca and P) | Ca | 2.44 | 2.09 | 1.74 | 1.39 | 1.04 | 0.70 | 0.35 | 0 |
|  | P | 0 | 0.31 | 0.62 | 0.93 | 1.24 | 1.55 | 1.86 | 2.17 |

#### 2.1.2. Image Acquisition

Images were acquired using a commercially-available spectral CT (MARS-12 v.5, MARS Bioimaging Ltd., Christchurch, NZ) equipped with a polychromatic X-ray source operating at 120 kVp with 2 mm aluminum beam filtration, and a PCD comprising three Medipix 3RX chips with CdZnTe semiconductor sensors utilizing five energy bins in charge summing mode (Ballabriga et al. 2013, Walsh et al. 2013). An evaluation of the X-ray attenuation profiles for elemental Ca and P (Berger *et al*. 2010, Hubbell and Seltzer 2004) revealed an intersection point at ∼44 keV when the thicknesses were selected to achieve equal attenuation across a 20-120 keV photon energy spectrum (Fig. 1). Energy thresholds were therefore selected to create five energy bins at 7-22, 22-32, 32-80, 80-90, and 90-120 keV, such that two bins were placed on either side of the Ca-P intersection point in the photon energy spectrum. These energy thresholds served to maximize differences in attenuation rate between Ca and P while maximizing signal from the lower energy thresholds. To mitigate the effects of pulse-pileup, the excitation current was set to 18 μA and each detector pixel was limited to a mean of 1000 total counts across all energy bins. Images were acquired using a helical scan with 1800 projections per rotation over two rotations and a 100 ms integration time. Reconstructions (MARS Bioimaging v1.1) were performed with a 200 μm isometric voxel size. For representative images, grayscale intensities were converted to Hounsfield units (HU) by internal calibration with air (−1000 HU) and water (0 HU).

**Figure 1.**
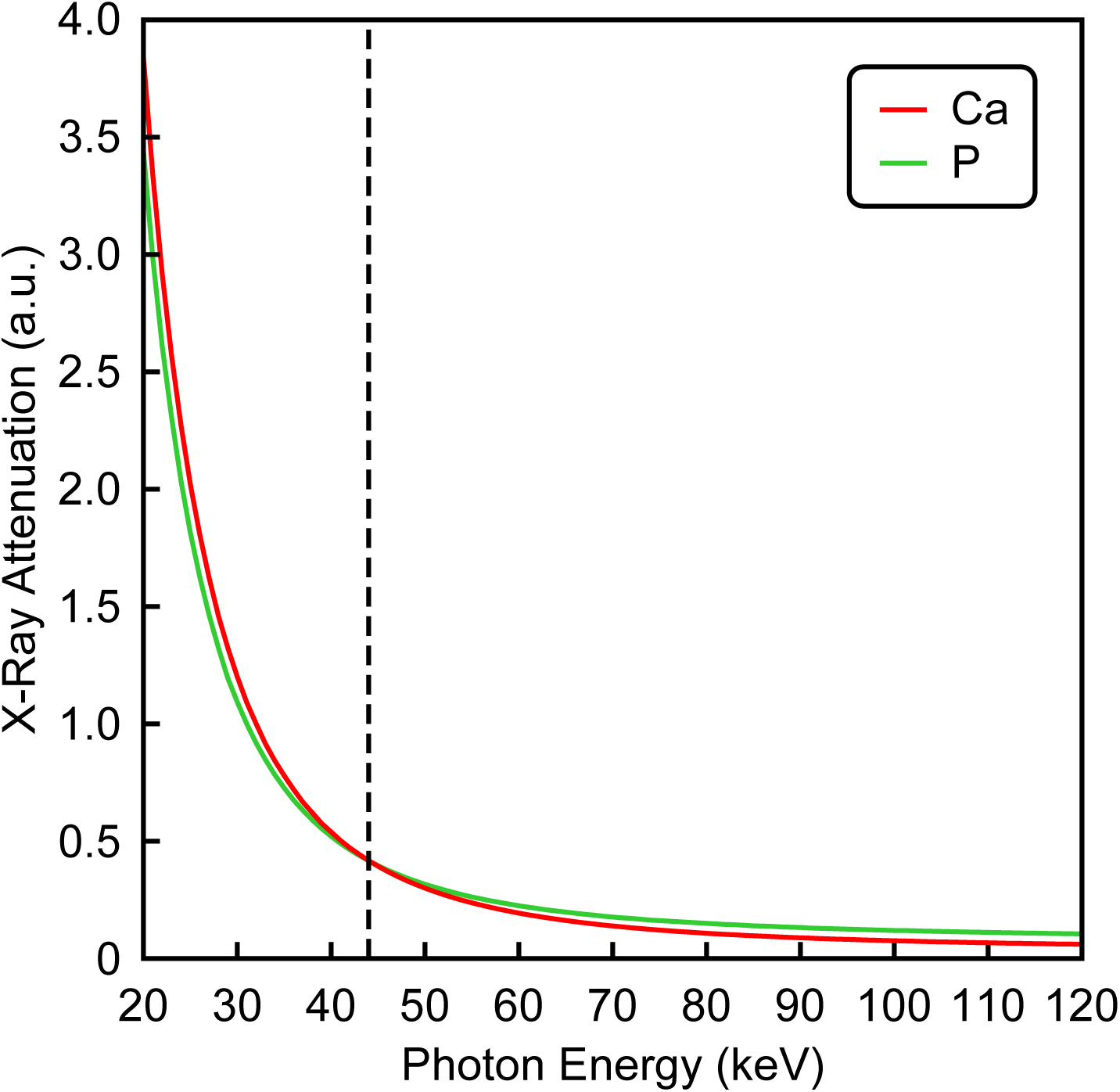
Calculated X-ray attenuation of Ca (red) and P (green) for a 120 kVp source with 2 mm aluminum filtration at thicknesses selected to achieve equivalent total attenuation across the 20-120 keV photon energy spectrum. An intersection point was observed at ∼44 keV where the energy dependent attenuation of P exceeds Ca at higher photon energies.

#### 2.1.3. Elemental Decomposition

Elemental decomposition of spatially coincident Ca, P, and water was performed using constrained maximum-likelihood estimation (MLE) in the image domain, adapting methods previously demonstrated for spatially discrete (Curtis and Roeder 2017) and coincident (Curtis and Roeder 2019) contrast and tissue compositions. Briefly, MLE was calibrated by a *M* x *N* material basis matrix, where *M* is the number of energy bins and *N* is the number of elements or materials to be decomposed. The material basis matrix was established by multiple linear least squares regression of the X-ray attenuation measured in each energy bin versus known concentrations of each material composition in the calibration phantom. The X-ray attenuation was measured as the mean attenuation within a 14.6 mm^3^ cuboidal volume of interest (VOI) located at the center of each Eppendorf tube in the calibration phantom images. Elemental volume fractions were estimated from the known concentrations of compositions in the calibration phantom based upon the mass and density of Ca, P, and water.

The volume fraction, or fractional abundance, of materials in each voxel of sample phantom images was determined by MLE using a quadratic programming function, quadprog, in MATLAB (v.9.0, Mathworks Inc.) such that solutions to the linear system of equations were constrained to full additivity and non-negativity. The estimated elemental volume fractions of unknown sample compositions were then scaled to millimolar (mM) concentration using the material basis matrix and linear regression model determined from the calibration phantom. The correlation coefficient (*R*^2^) between the measured attenuation values used for calibration of the elemental decomposition of Ca and P across all energy bins was calculated by linear least square regression using MATLAB (v.9.0, Mathworks Inc.).

The accuracy of material decomposition was evaluated from sample phantom images using a 17.6 mm^3^ cuboidal VOI, including 40 slices of the material decomposition, located at the center of each Eppendorf tube. Sensitivity and specificity for detecting Ca and P were calculated using the results from the mixed composition solutions. Mixtures containing 1.04 M (41.7 g/L) Ca and 1.24 M (38.4 g/L) P were selected as true positive VOIs (10 total) for each element. An equal total number of true negative VOIs (10 total) were selected as those containing 0 mM of the element of interest. AUC was calculated by evaluating sensitivity and specificity over a range of concentration thresholds from 0 to 5 M in 0.01 M increments, which exceeded the concentrations within the sample phantom, to determine sets of positive and negative voxels. Sensitivity and specificity were reported at a 0.5 M threshold, which was approximately 20% of the maximum concentration in any well. The quantitative accuracy of elemental decomposition was evaluated by the root mean squared error (RMSE) of all estimated versus known concentrations for Ca and P.

### 2.2. Calcification Model

Elemental decomposition of Ca, P, and O in clinically-relevant μcal compositions was investigated with model μcals containing varying ratios, or mole fractions, of Ca and P (Table 2). The percentage contribution of each elemental component in HA and CaOx to the overall mass and X-ray attenuation was calculated based upon the mole fractions, molecular weights, and NIST attenuation values (Berger *et al*. 2010, Hubbell and Seltzer 2004) integrated over the photon energy spectrum of the tube emission measured by the PCD (9-120 keV) (Table 2). HA and CaOx exhibit a P:Ca ratio of 3:5 and 0:1, respectively. Therefore, DCP (P:Ca = 1:1) was added as a third μcal composition, and as a positive-control for P, to investigate the ability to decouple the contribution of each element from the overall attenuation of the μcal composition.

**Table 2.** Percent contribution of each element to the overall moles, mass, and X-ray attenuation of each μcal composition.

| | μcal | Composition | Ca | P | O | H | C |
| --- | --- | --- | --- | --- | --- | --- | --- |
| Mole<br>Percent | CaOx | $\text{CaC}_2\text{O}_4$ | 14 | 0 | 57 | 0 | 29 |
| | HA | $\text{Ca}_5(\text{PO}_4)_3\text{OH}$ | 23 | 14 | 59 | 5 | 0 |
| | DCP | $\text{CaHPO}_4$ | 14 | 14 | 57 | 14 | 0 |
| Mass<br>Percent | CaOx | $\text{CaC}_2\text{O}_4$ | 24 | 0 | 59 | 2 | 15 |
|  |  |  | 31 | 0 | 50 | 0 | 19 |
| | HA | $\text{Ca}_5(\text{PO}_4)_3\text{OH}$ | 40 | 18 | 41 | 0 | 0 |
| | DCP | $\text{CaHPO}_4$ | 29 | 23 | 47 | 1 | 0 |
| Attenuation<br>Percent | CaOx | $\text{CaC}_2\text{O}_4$ | 82 | 0 | 16 | 0 | 2 |
| | HA | $\text{Ca}_5(\text{PO}_4)_3\text{OH}$ | 78 | 16 | 7 | 0 | 0 |
| | DCP | $\text{CaHPO}_4$ | 68 | 23 | 9 | 0 | 0 |

#### 2.2.1. Calibration and Sample Phantom

Model μcals comprised discrete concentrations of HA, CaOx, and DCP placed within a modular phantom. Biomimetic HA, Ca_5_(PO_4_)OH, crystals were prepared using previously established methods (Roeder *et al*. 2006). Suspensions of HA or commercially-available DCP, CaHPO_4_·2H_2_O (Fisher Scientific), crystals were prepared in DI water containing 1% agarose at a concentration of 0, 3.75, and 7.5 vol%. Suspensions of commercially-available CaOx, CaC_2_O_4_·H_2_O (Sigma Aldrich, ≥ 99.9%), crystals were also prepared in DI water containing 1% agarose at a concentration of 0, 7.5, and 11.25 vol%. The concentrations of CaOx were chosen to match the total attenuation of HA across the overall photon energy spectrum of the polychromatic X-ray source (∼20-120 keV). Eppendorf tubes (5 mm inner diameter) containing each discrete composition and concentration (7 total) were placed in a modular acrylic phantom (35 mm outer diameter) for imaging in a single acquisition. A polytetrafluoroethylene screw was present at the center of the phantom as part of a clamping mechanism for holding Eppendorf tubes within the phantom and was excluded from material decomposition. The same phantom was used for both calibration and evaluation.

#### 2.2.2. Image Acquisition

Images were acquired using a commercially-available spectral CT (MARS-12 v.5, MARS Bioimaging Ltd., Christchurch, NZ) equipped with a polychromatic X-ray source operating at 120 kVp with 0.2 mm copper beam filtration, and a PCD comprising three Medipix 3RX chips with CdZnTe semiconductor sensors utilizing five energy bins in charge summing mode (Ballabriga et al. 2013, Walsh et al. 2013). Energy thresholds were selected to create five energy bins at 7-25, 25-35, 35-74, 74-87, and 87-120 keV, such that two bins were placed on either side of the Ca-P intersection point in the photon energy spectrum (Fig. 1). These energy thresholds served to maximize differences in attenuation rate between Ca and P while maximizing signal from the lower energy thresholds and ensuring adequate count statistics in each energy bin (≥ 100 counts) for image reconstruction. To mitigate the effects of pulse-pileup, the excitation current was set to 18 μA and each detector pixel was limited to a mean of 1000 total counts across all energy bins. Images were acquired using a circular scan with 1800 projections per rotation over two rotations and a 100 ms integration time. Reconstructions (MARS Bioimaging v1.1) were performed with a 200 μm isometric voxel size. For representative images, grayscale intensities were converted to Hounsfield units (HU) by internal calibration with air (−1000 HU) and water (0 HU).

#### 2.2.3. Elemental Decomposition

Elemental decomposition of Ca, P, O, and water was performed using constrained MLE in the image domain. Methods previously developed for decomposition of spatially coincident materials were adapted to allow calibration from multiple elements within the same voxel (Curtis and Roeder 2019). Given the known concentration of the μcal composition (HA, CaOX, and DCP) in model

μcals, the theoretical volume fraction, *V’*, of each μcal composition, *j*, and water in model μcals was calculated as,

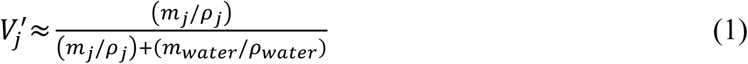

where *j* is the μcal composition (HA, CaOX, and DCP), *m*_*j*_ and *m*_*wate*r_ are the mass of each μcal composition and water, respectively, *ρ*_*j*_ and *ρ*_*water*_ are the density of each μcal composition and water, respectively, and the volume fraction of water in the model μcal is *V′*_*water*_ ≈ 1 – *V′*_*j*_.

The elemental volume fraction was calculated using two methods. In the first approach, each atom within the crystal structure of each μcal composition was assumed to have equal volume, producing an elemental isovolumetric decomposition. The molar fraction, *M*, of each element, *i*, within each μcal composition, *j*, was calculated as,

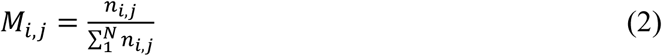

where *n*_*i,j*_ is the number of moles of element, *i*, present in μcal composition, *j*, and *N* is the total number of elements (Ca, P, and O). The volume fraction, *V’*, of each element, *i*, in each μcal ccomposition, *j*, in model μcals was subsequently estimated as,

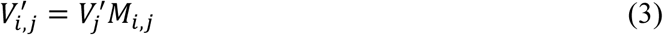

where *V′*_*j*_ is the volume fraction of the μcal composition, *j*, in the model μcal calculated from equation (1) and *M*_*i,j*_ is the molar fraction of each element, *i*, within each μcal composition, *j*, calculated from equation (2).

In the second approach, the true volume fraction of each element within each μcal composition was estimated using atomic radii. The total volume, *v*, of each element, *i*, in each μcal composition, *j*, was estimated as,

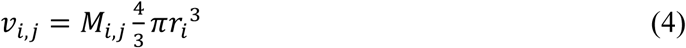

where *M*_*i,j*_ is the molar fraction of each element, *i*, within each μcal composition, *j*, calculated from equation (2) and *r* is the atomic radius for each element, *i*. Atomic radii were utilized for both ionic bonding of Ca (1.26 Å), P (0.58 Å), and O (1.28 Å), and covalent bonding of Ca (1.74 Å), P (1.06 Å), and O (0.73 Å) (Cordero *et al*. 2008). The volume fraction, *V*, of each element, *i*, within each μcal composition, *j*, was estimated as,

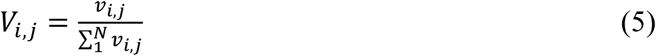

where *v*_*i,j*_ is the total volume of each element, *i*, within each μcal composition, *j*, calculated from equation (4) and *N* is the total number of elements (Ca, P, and O). The volume fraction, *V’*, of each element, *i*, for each μncal composition, *j*, in model μcals was estimated as,

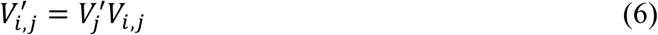

where *V′*_*j*_ is the volume fraction of the μcal composition, *j*, in the model μcal calculated from equation (1) and *V*_*i,j*_ is the volume fraction of each element, *i*, within each μcal composition, *j*, calculated from equation (5). Water was considered as a wholly separate endgroup for material decomposition. Note that elemental O and water (H_2_O) are able to be decomposed as separate endgroups by MLE because the density and attenuation of elemental O is much greater than that of water.

Elemental decomposition was performed on the same images used for calibration. For each method of elemental decomposition, the volume fraction contributions of Ca, P and O were averaged across a 20 slice volume, assigned to RGB channels, respectively, and mapped for model μcals. The mole fraction of P, P/(Ca+P), was also mapped to discriminate all three μcal compositions. Images were scaled from 0 to 1.25 times the maximum volume fraction of each element in the calibration matrix. This ensured that brightness and contrast within each image was representative of the overall range for each elemental calibration.

The accuracy of elemental decomposition, without scaling, was evaluated for elemental isovolume and true volume decompositions. The sensitivity and specificity for discriminating one μcal composition from another was calculated for elemental Ca, P, and O concentrations, as well as the mole fraction of P. Model μcals containing two concentrations of the μcal composition of interest were selected as true positive VOIs for each element. An equal total number of true negative VOIs were selected from two concentrations of the other μcal compositions. For notation, the true positive μcal composition of interest is hereafter listed before the negative μcal composition (e.g., HA versus CaOx). AUC was calculated by evaluating sensitivity and specificity over a range of thresholds including 0 to 2 times the maximum value measured in the μcal composition of interest with 400 equidistant threshold steps, to determine sets of positive and negative voxels. The dimensions (diameter and length) of the evaluation VOI was varied to determine detection limits for accurate μcal identification and to evaluate size-dependent discrimination of the μcal compositions (Table 3 and Fig. 2). The size of the VOI was varied to include and exceed the size of pathologically-relevant μcals. Measured AUC values across all elements and VOIs were compared between calibration methods using a paired *t*-test (MATLAB). The quantitative accuracy of elemental decomposition was evaluated by the root mean squared error (RMSE) of all estimated versus known concentrations for each element using the mean volume fraction values measured from 15.02 mm^3^ cylindrical VOIs at the center of each sample (*n* = 7).

**Table 3.** Variation in the VOI dimensions used for evaluating the accuracy of elemental decomposition.

| ID | Dimension Limit (mm) | Volume (mm<sup>3</sup>) | Number of VOIs | Total Volume (mm<sup>3</sup>) |
| --- | --- | --- | --- | --- |
| 0 | 0.20 | 0.01 | 80 | 0.64 |
| 1 | 0.60 | 0.17 | 40 | 6.79 |
| 2 | 1.00 | 0.79 | 20 | 15.71 |
| 3 | 1.40 | 2.16 | 12 | 25.86 |
| 4 | 1.80 | 4.58 | 8 | 36.64 |
| 5 | 2.20 | 8.36 | 8 | 66.90 |
| 6 | 2.60 | 13.80 | 4 | 55.22 |
| 7 | 3.00 | 21.21 | 4 | 84.82 |
| 8 | 3.40 | 30.87 | 4 | 123.48 |
| 9 | 3.80 | 43.10 | 4 | 172.39 |
| 10 | 4.20 | 58.19 | 4 | 232.75 |

**Figure 2.**
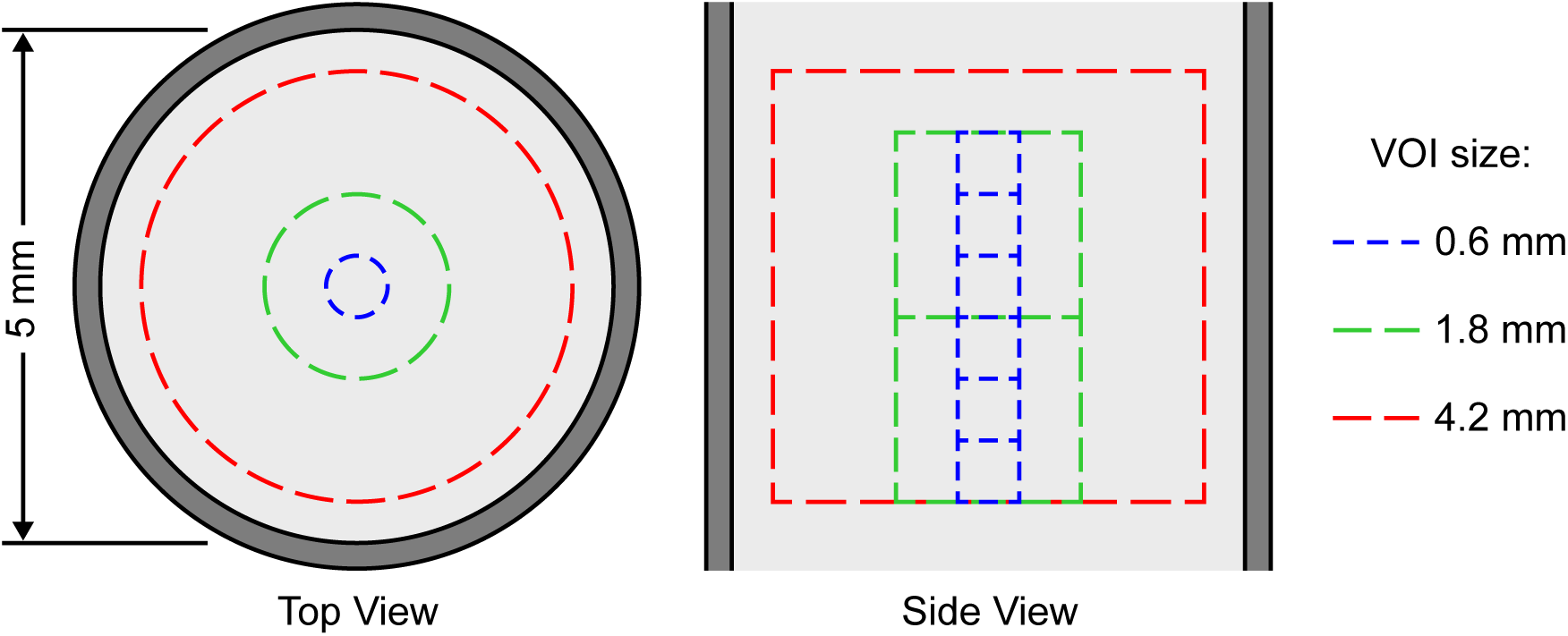
Schematic diagram of three example VOIs confined to a maximum dimension (0.6, 1.8 and 4.2 mm diameter and length) showing the volume and number of replicates available within the overall sample volume.

## 3. Results and Discussion

### 3.1. Homogenous Liquid Phantom

Photon-counting spectral CT enabled elemental decomposition of P and Ca in mixed, or spatially coincident, compositions of the liquid phantom (Fig. 3) The correlation coefficient of attenuation curves for both Ca and P was 0.997. Nonetheless, small differences in the attenuation curves permitted elemental decomposition (Fig. 3). The mean AUC, measured for both Ca and P was 0.80, which indicated accurate discrimination of Ca and P in mixed compositions. The RMSE of measured concentrations was 0.17 and 0.22 M for Ca and P, respectively, which was 6.7% of the maximum Ca concentration and 10.1% of the maximum P concentration (Fig. 4). Importantly, the RMSE as a percent of the maximum signal for measurement of Ca and P concentrations was comparable in magnitude to that previously measured for discrete contrast agents (Curtis and Roeder 2017).

**Figure 3.**
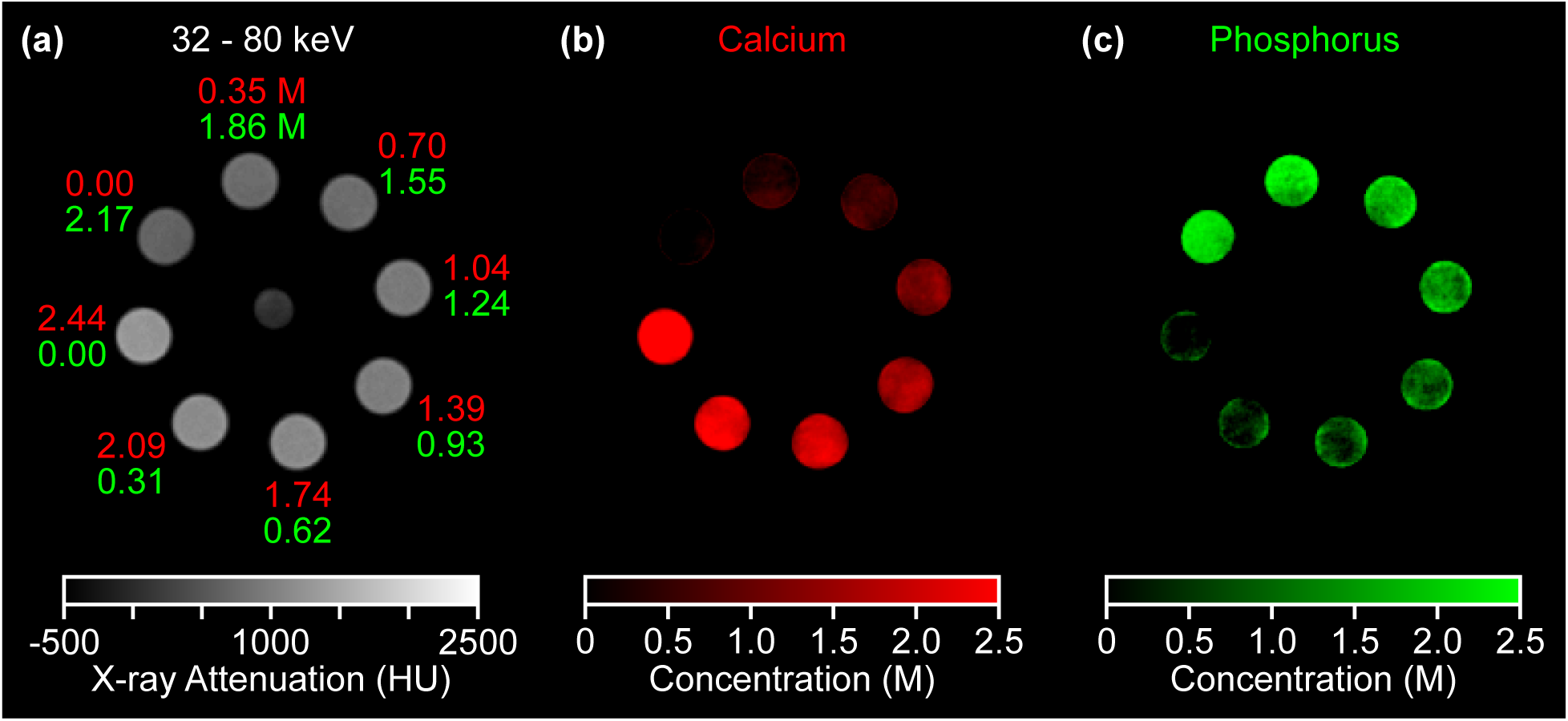
Representative image slices of a sample phantom with mixed, or spatially coincident, compositions comprising calcium chloride ranging from 0 to 2.44 M and phosphoric acid ranging from 0 to 2.17 M showing (a) grayscale image from the middle energy bin (32-80 keV) and quantitative material decomposition maps of (b) Ca (red) and (c) P (green). Grayscale intensities were converted to HU and color intensities were scaled to molar concentrations.

**Figure 4.**
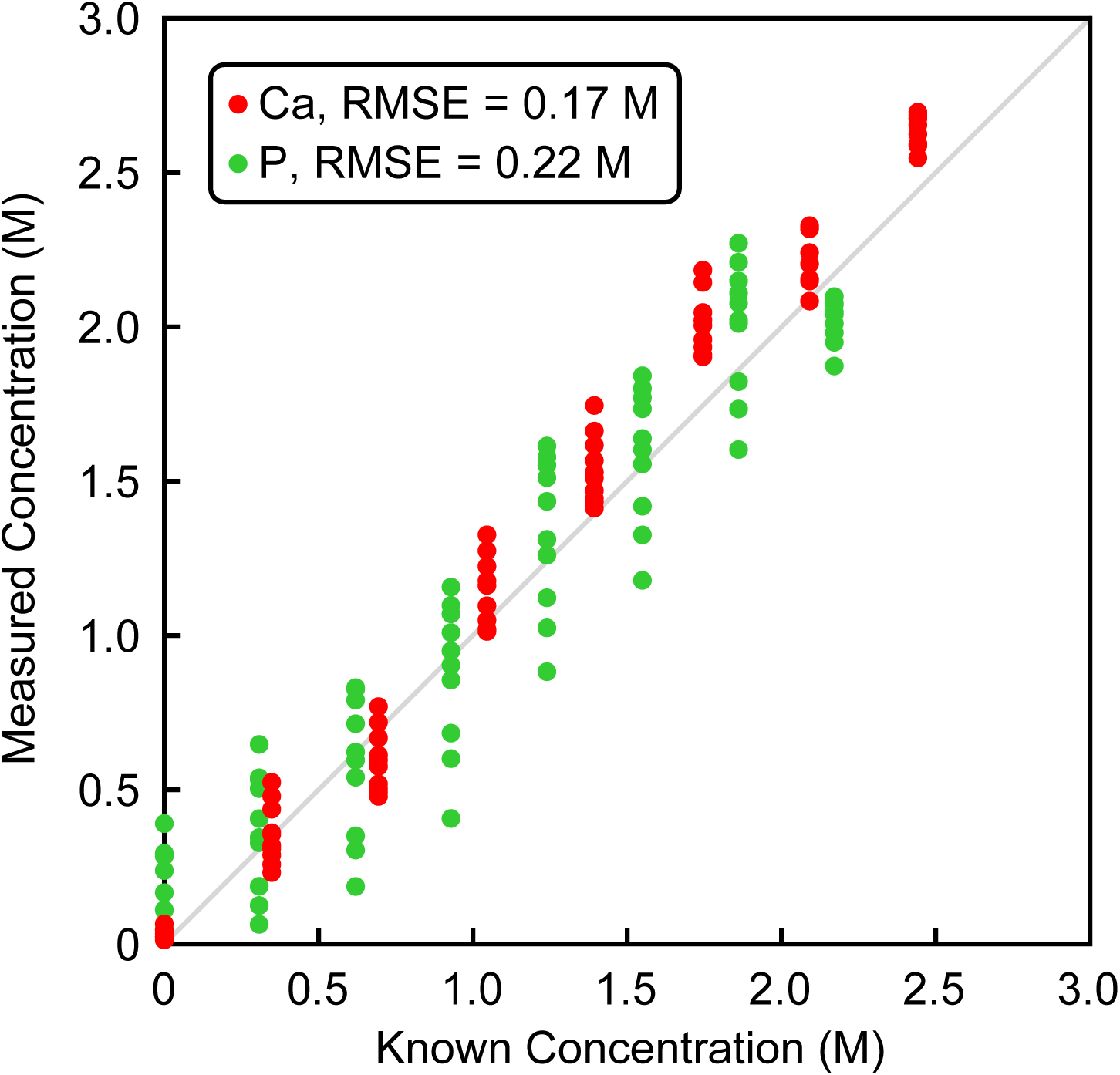
Linear regression of measured versus known concentrations of Ca (red) and P (green) in mixed compositions of the homogenous liquid phantom.

### 3.2. Calcification Model

Adjustments were made in the overall protocol to mitigate reconstruction artifacts that could be caused by low count rates at the detector. A 0.2 mm copper filter was used in place of the 2 mm aluminum filter in order to reduce beam hardening artifacts in the lowest bins. Bin widths were also adjusted to have a minimum of 100 counts per bin to ensure count statistics were adequate for reconstruction. The highest thresholds were reduced to enhance counts in the upper bins leading to thresholds of 25, 35, 74 and 87 keV. Image projections were acquired using a circular scan. All other parameters remained the same as the previous experiment.

The fractional contribution of each element to the overall attenuation of each crystal composition revealed a significant contribution from oxygen (Table 2). Initial attempts to apply the Ca and P elemental decomposition from the homogenous liquid phantoms to the calcification models using matching protocol settings resulted in an erroneously high attribution of P to CaOx wells. The attenuation from oxygen, which was not accounted for in the elemental decomposition, was erroneously identified as P. Thus, the inclusion of oxygen as a separate element in the calcium crystal compositions proved vital to achieving an accurate elemental decomposition.

Material decomposition using the elemental isovolume, ionic true volume, or the covalent true volume assumption enabled quantitative elemental decomposition of Ca, P and O within each calcium crystal composition (Figs. 5-7). Qualitatively a clear delineation between Type I (CaOx) and Type II (HA) μcals was observed based on the P content (Figs. 5d, 6d, and 7d). Moreover, mapping the P/(Ca+P) ratio qualitatively showed clear delineation between Type I and Type II μcals irrespective of the concentration observed within each well (Figs. 5e, 6e, and 7e). This demonstrated that Type I (CaOx) and Type II (HA) μcals were able to be discriminated despite changes in their overall attenuation due to concentration (density). Elemental composition maps produced using the isovolume and ionic true volume assumptions were nearly identical.

**Figure 5.**
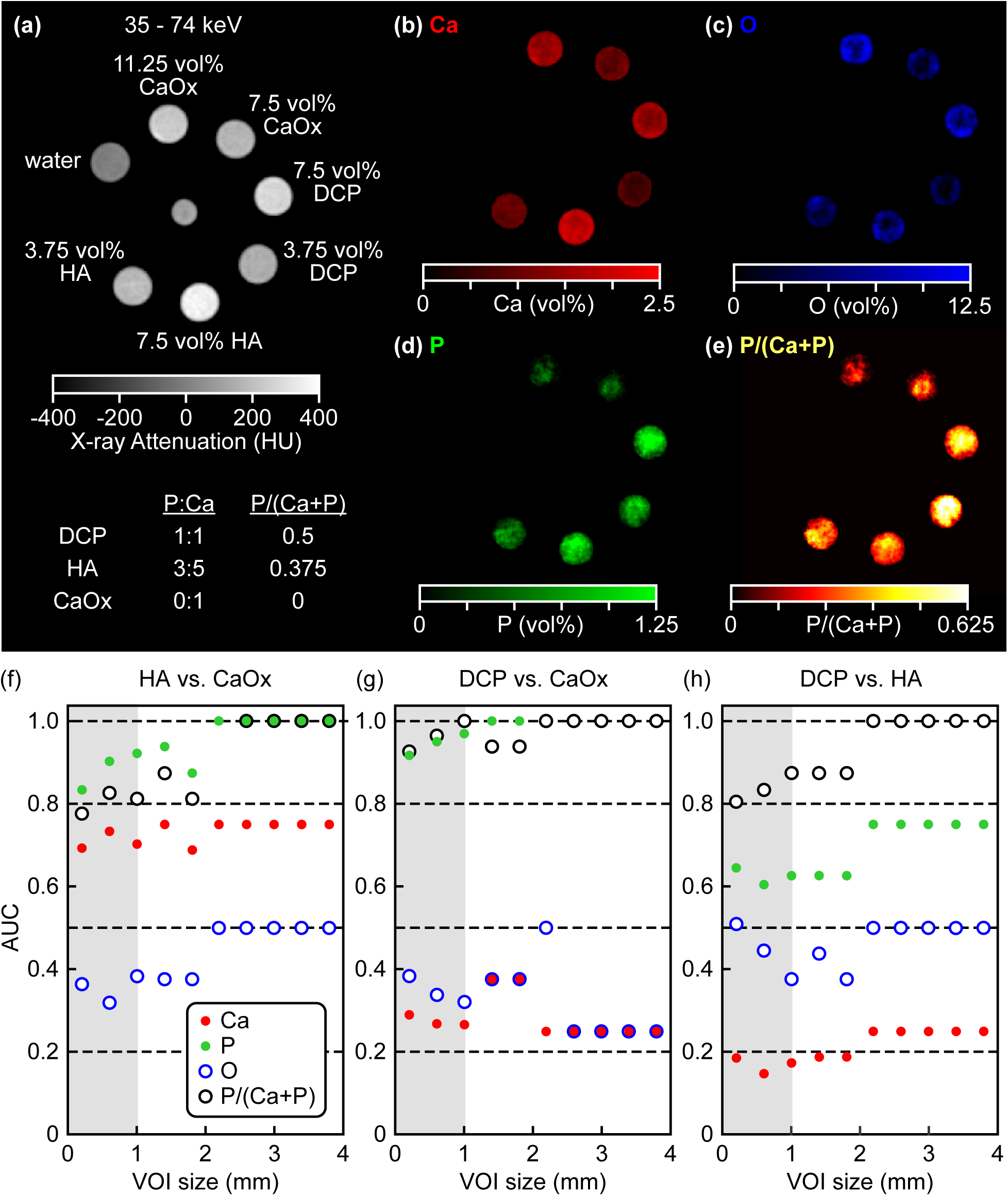
Representative image slices of a sample phantom with model μcals comprising HA and DCP ranging in concentration from 0 to 7.5 volume percent, and CaOx ranging from 0 to 11.25 volume percent, including a (a) grayscale image from the middle energy bin (32-74 keV) and quantitative elemental decomposition maps using the elemental isovolumetric calibration showing (b) Ca (red), (c) O (blue), (d) P (green) and (e) the P/(Ca+P) ratio. Grayscale intensities were converted to HU and color intensities were scaled to volume percent or the P/(Ca+P) ratio.

Type I μcals (CaOx) lack P. Thus, differences in the measured P content of μcals provides a strong positive radiographic marker for identifying HA versus CaOx. False P signal was present in the CaOx crystal composition for both the elemental true volume and elemental isovolume assumptions, albeit at a much lower concentration than the P concentration observed in HA μcals. Moreover, the false signal from P in CaOx μcals was lower using the covalent true volume fraction calibration (Fig. 7) compared with either the elemental isovolume or the ionic true volume fractions (Figs. 5 and 6). This result suggests that the covalent true volume decomposition was able to more closely approximate the elemental composition of the calcium crystal compositions, most likely because the percent contribution of each element to the overall volume fraction (Table 4) and X-ray attenuation (Table 2) were more closely matched for the assumption of covalent true volume compared with either elemental isovolume or ionic true volume (Table 5).

**Table 4.**
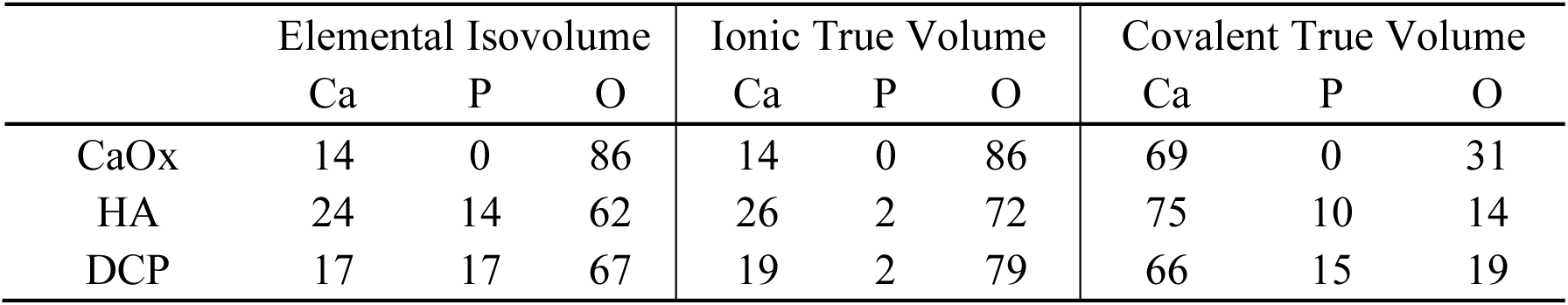
Percent contribution of each element to the overall volume fraction of each calcium crystal composition.

**Table 5.**
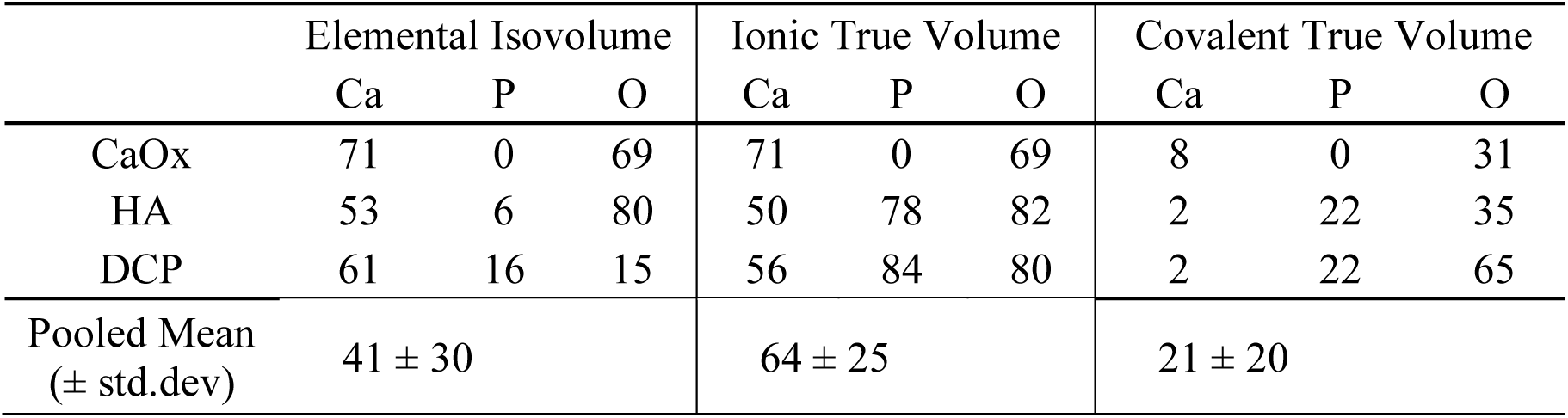
Percent difference between the contribution of each element to the total X-ray attenuation contribution and volume fraction of each calcium crystal composition.

**Figure 6.**
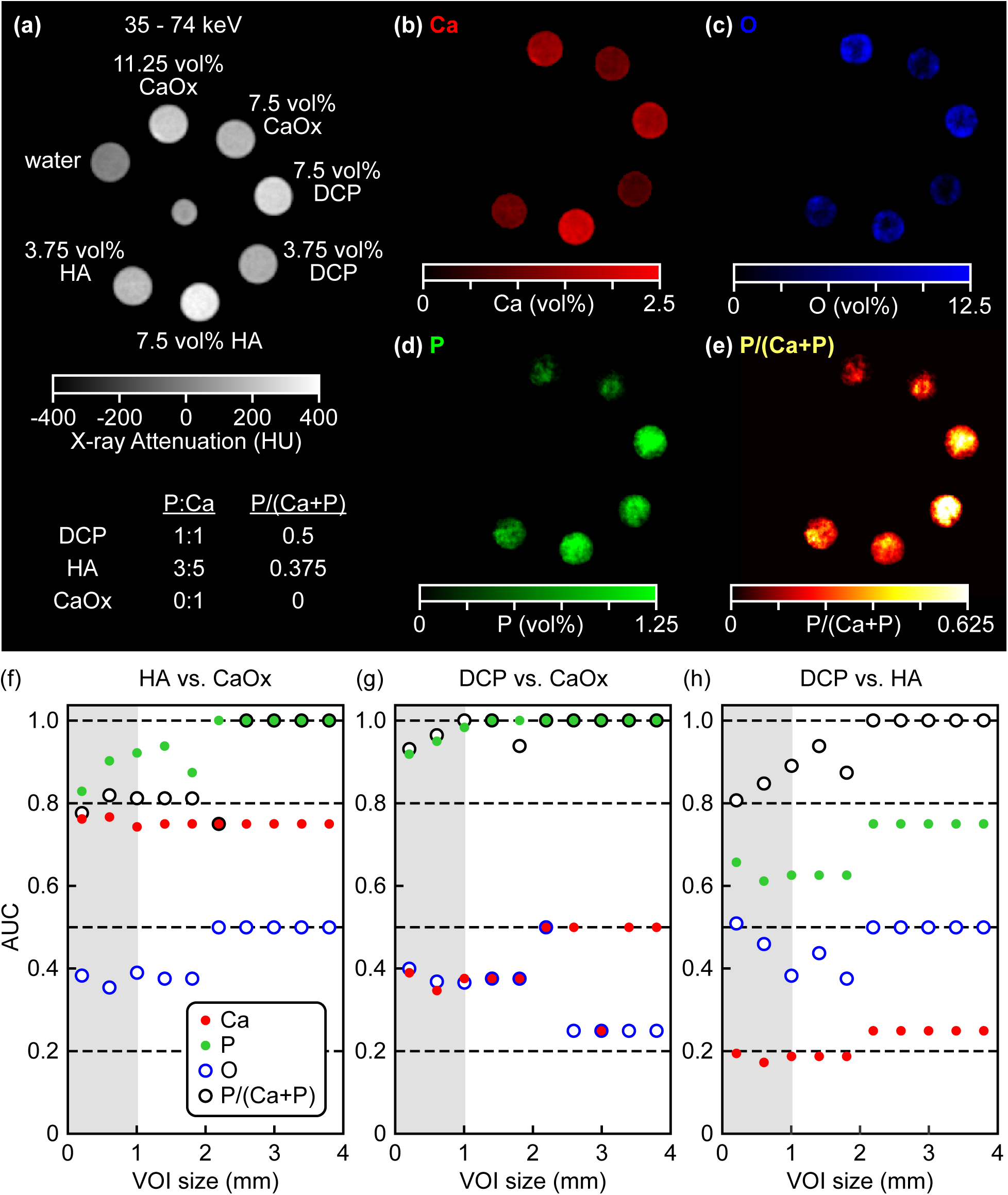
Representative image slices of a sample phantom with model μcals comprising HA and DCP ranging in concentration from 0 to 7.5 volume percent, and CaOx ranging from 0 to 11.25 volume percent, including a (a) grayscale image from the middle energy bin (32-74 keV) and quantitative elemental decomposition maps using the ionic true volume fraction calibration showing (b) Ca (red), (c) O (blue), (d) P (green) and (e) the P/(Ca+P) ratio. Grayscale intensities were converted to HU and color intensities were scaled to volume percent or the P/(Ca+P) ratio.

**Figure 7.**
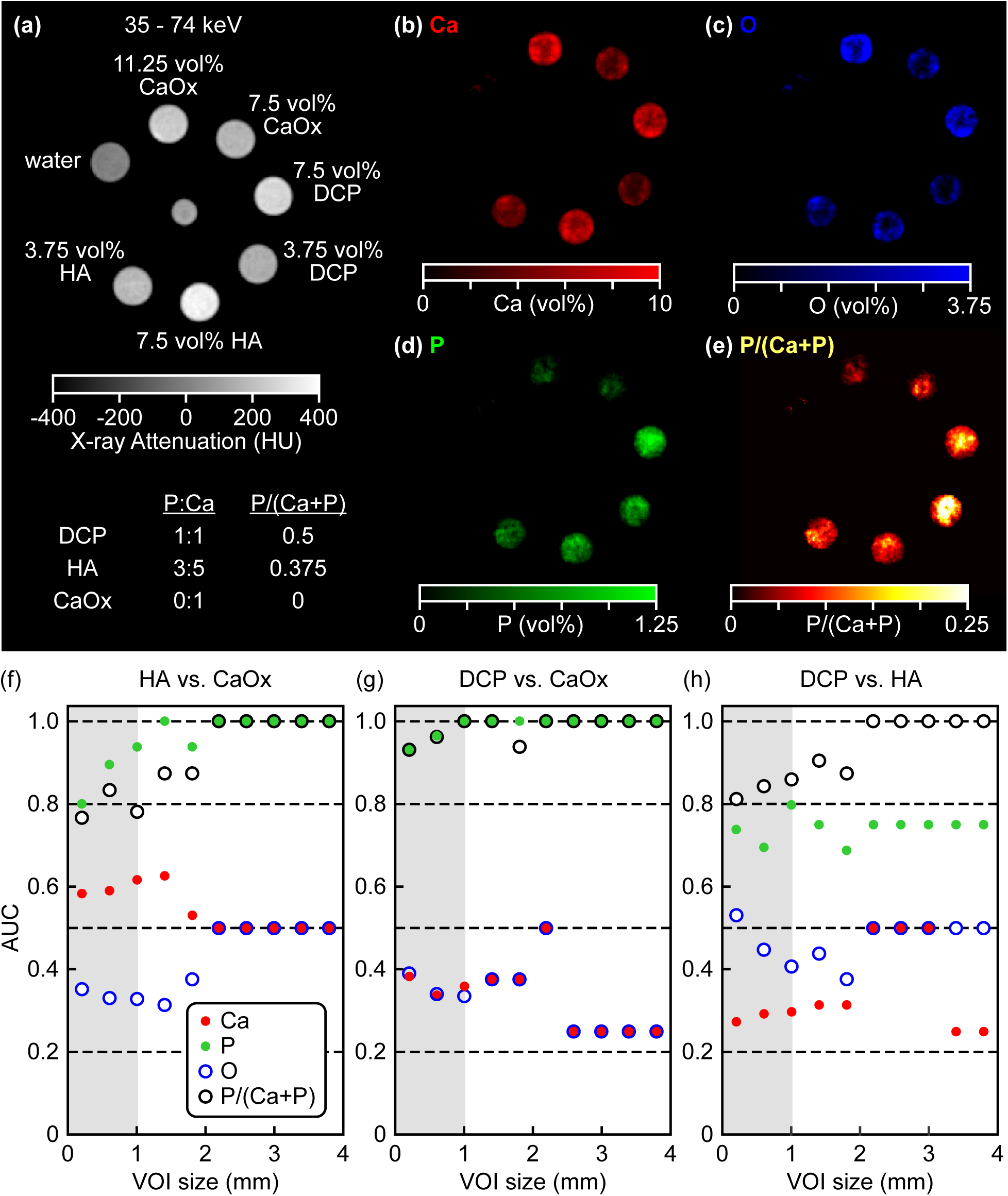
Representative image slices of a sample phantom with model μcals comprising HA and DCP ranging in concentration from 0 to 7.5 volume percent, and CaOx ranging from 0 to 11.25 volume percent, including a (a) grayscale image from the middle energy bin (32-74 keV) and quantitative elemental decomposition maps using the covalent true volume fraction calibration showing (b) Ca (red), (c) O (blue), (d) P (green) and (e) the P/(Ca+P) ratio. Grayscale intensities were converted to HU and color intensities were scaled to volume percent or the P/(Ca+P) ratio.

**Figure 8.** Quantitative analysis of elemental isovolume decomposition showing the AUC for delineation of (a) HA versus CaOx, (b) DCP versus CaOx, and (c) DCP versus HA calcium crystal compositions, using the measured elemental concentrations of Ca (red), P (green), O (blue), and P/(Ca+P) (purple), within a VOI of varying size (maximum dimension). The shaded region shows clinically relevant μcal dimensions. Dashed lines show AUC levels that provide a strong indicator for the positive group (green) (e.g., HA in HA versus CaOx) and negative group (red) (e.g., CaOx in HA versus CaOx). A measured AUC of 0.5 (gray dashed line) indicates no ability to distinguish between two compositions.

**Figure 9.** Quantitative analysis of ionic true volume decomposition showing the AUC for delineation of (a) HA versus CaOx, (b) DCP versus CaOx, and (c) DCP versus HA calcium crystal compositions, using the measured elemental concentrations of Ca (red), P (green), O (blue), and P/(Ca+P) (purple), within a VOI of varying size (maximum dimension). The shaded region shows clinically relevant μcal dimensions. Dashed lines show AUC levels that provide a strong indicator for the positive group (green) (e.g., HA in HA versus CaOx) and negative group (red) (e.g., CaOx in HA versus CaOx). A measured AUC of 0.5 (gray dashed line) indicates no ability to distinguish between two compositions.

**Figure 10.** Quantitative analysis of covalent true volume decomposition showing the AUC for delineation of (a) HA versus CaOx, (b) DCP versus CaOx, and (c) DCP versus HA calcium crystal compositions, using the measured elemental concentrations of Ca (red), P (green), O (blue), and P/(Ca+P) (purple), within a VOI of varying size (maximum dimension). The shaded region shows clinically relevant μcal dimensions. Dashed lines show AUC levels that provide a strong indicator for the positive group (green) (e.g., HA in HA versus CaOx) and negative group (red) (e.g., CaOx in HA versus CaOx). A measured AUC of 0.5 (gray dashed line) indicates no ability to distinguish between two compositions.

Quantitative evaluation of the accuracy of elemental decomposition using the elemental isovolume and covalent true volume assumptions revealed a low quantitative error (RMSE) for all elements when measured against expected values for each element and concentration (Table 6). The greatest deviations were observed in measured oxygen values due to low overall attenuation from the element (Table 2). Coefficients of determination (*R*^2^) above 0.7 for Ca and P were achieved across a 1:1 regression of the dataset onto the calibration matrix. The quantitative accuracy (*R*^2^ and RMSE) of P content did not appear to be significantly affected by the assumption of the elemental decomposition (Table 6). This result suggests that the quantitative elemental decomposition was robust against large fluctuations in volume fraction values assigned for calibration (Table 4). RMSE was increased and *R*^2^ decreased for Ca, with an opposite effect observed for oxygen when the covalent true volume calibration was compared to the elemental isovolume calibration (Table 6). The estimated contribution of Ca to the overall volume of each calcium crystal composition was lower within the covalent true volume calibration (Table 4). Thus, the variability of the attenuation within each model μcal likely had a greater impact on the quantitative accuracy of the covalent true volume decomposition compared with the elemental isovolume decomposition. However, reconstruction artifacts were not accounted for in this analysis and additional imaging replicates are required to confirm this interpretation.

**Table 6.**
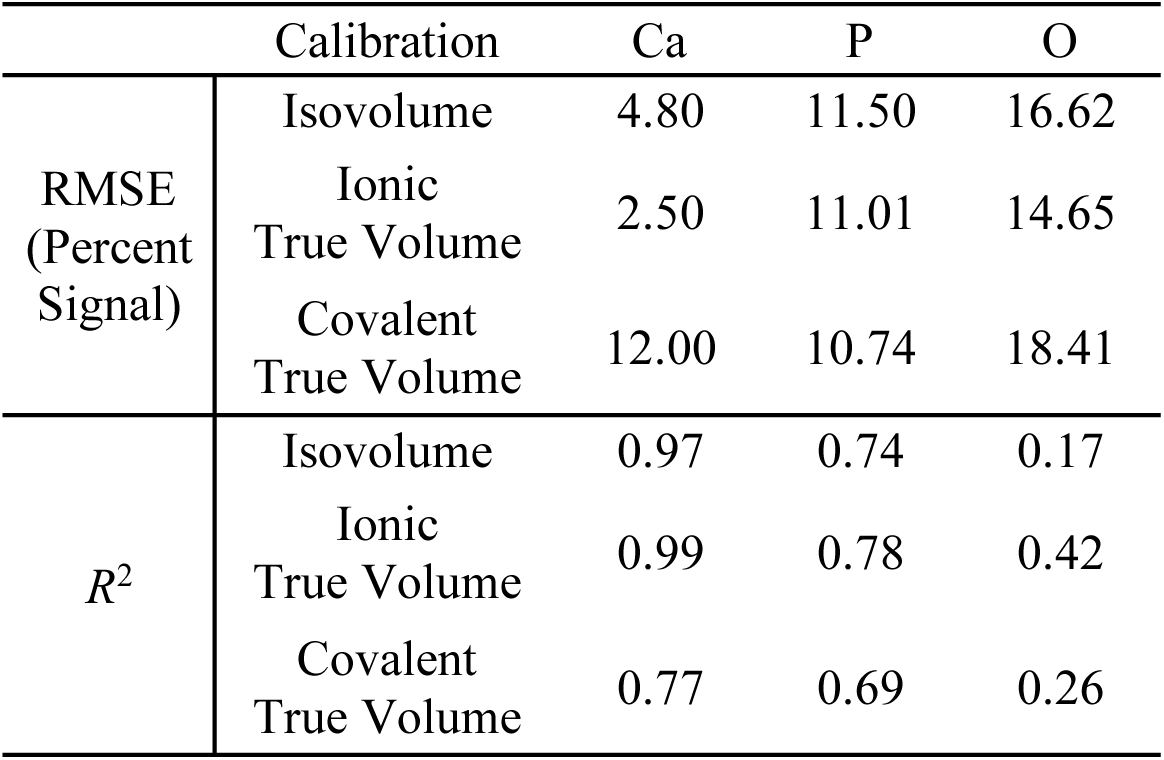
RMSE and coefficient of determination for each element using the elemental isovolume and true volume assumptions in the elemental decomposition.

Quantitative evaluation of the effect of the VOI on AUC revealed common trends between the elemental isovolume decomposition (Fig. 5), the ionic true volume decomposition (Fig. 6) and the covalent true volume decomposition (Fig. 7). In all decompositions, oxygen content was ineffective (AUC = 0.5) for the delineation of HA from either CaOx or DCP, and exhibited moderate ability to delineate CaOx from DCP. The inability of the oxygen content to accurately separate DCP and HA was due to only a 2% difference in the fractional contribution of O to the overall attenuation of DCP and HA (Table 2). In preliminary studies, cross-contamination was observed in the measurement of P and O. Thus, the higher contribution of P to the DCP signal may have aided in the delineation of DCP from CaOx using measurements of the oxygen content.

Importantly, the P/(Ca+P) ratio was the only metric that provided a significant positive indicator (AUC > 0.8) for the delineation of all three calcium crystal compositions, regardless of the calibration method used. This was due, in part, to the concentration independency of the P/(Ca+P) ratio. All other metrics required the exact concentration of the element within the crystal structure to be evaluated using AUC. Multiple concentrations of each calcium crystal composition were used for analysis and lower concentrations of each crystal were less likely to surpass the thresholds selected for evaluation of specificity and sensitivity, causing AUC to fall below 0.8. The only exception to this was the comparison of P-containing crystal compositions, DCP and HA, to CaOx which lacks P, using the measured P content. In this case, the measured signal from P in CaOx μcals was merely a consistent background level of erroneous signal (noise) which was substantially lower than the P content measured for either DCP or HA μcals (Figs. 5 and 7). Consequently, all concentrations of CaOx were able to be accurately thresholded from all concentrations of DCP and HA using the P content for all assumptions used in the elemental decomposition. Thus, the use of the P content for analysis in the P/(Ca+P) ratio enabled accurate discrimination of all three calcium crystal compositions using all elemental decomposition calibrations.

The AUC for elemental isovolume, ionic true volume, and covalent true volume decompositions exhibited the greatest variability when HA was included in the analysis (Figs. 5-7). DCP and CaOx presented the largest differences in P content (Table 2) and were therefore more readily identified using P content and P/(Ca+P). HA exhibits P content in between that of CaOx and DCP, and was therefore the most difficult to separate from either DCP or CaOx. Thus, identification of HA was more dependent on the measured Ca content. The discrimination of HA versus CaOx or DCP using the measured Ca from the elemental isovolume and ionic true volume assumptions trended towards a distinct separation (AUC=0.8 and 0.2, respectively) (Figs. 5 and 6), while the same evaluation using the covalent true volume assumption trended towards an inability to discriminate HA (AUC=0.5) (Fig. 7). The percent contribution of Ca to both elemental isovolume fraction and ionic volume fraction exhibited approximately a 30 and 52 percent difference for HA versus DCP and CaOx, respectively (Table 4). Conversely, the covalent true volume fractional contribution of Ca exhibited an 8 and 12 percent difference for HA versus CaOx and DCP, respectively (Table 4). Therefore, the elemental isovolume and ionic true volume allowed a greater relative dynamic range for evaluating Ca volume fraction between the calcium crystal compositions, allowing more discrete thresholding of the Ca volume fraction for discriminating the HA μcals. Notably though, a paired *t*-test of the AUC results comparing each elemental decomposition revealed that there were no significant differences in AUC across all calcium crystal composition pairs (*p* = 0.86). Thus, major variations in the calibration did not significantly impact the overall capability of the image-based elemental decomposition to perform classification tasks for each calcium crystal composition. This result suggests that the image-based elemental decomposition was robust to fluctuations in the volume fractions of the calibration used. Therefore, elemental calibration can be readily implemented for discrimination of calcium crystal compositions.

Finally, to be readily applicable in a clinical setting Type I (CaOx) and Type II (HA) μcals must be able to be discriminated at a maximum dimension not exceeding 1 mm with an AUC > 0.8. AUC for the P/(Ca + P) ratio exceeded 0.8 within this clinical size range (Figs. 5-7). Therefore, the P/(Ca+P) ratio was able to discriminate a variety of calcium crystal compositions, independent of their concentration. However, the elemental volume fraction of P provided stronger discrimination of Type I (CaOx) and Type II (HA) μcals at the smallest VOI analyzed, (AUC ≥ 0.8), because CaOx lacks P content (Table 2, Figs. 5-7). Therefore, the P content is recommended for absolute discrimination of Type I versus Type II μcals, while the P/(Ca+P) ratio can be more generally applied to identify various calcium crystal compositions containing distinct P:Ca ratios. Classification of HA and CaOx was investigated previously using the MARS Bioimaging photon-counting spectral CT (Kirkbride *et al*. 2017). The attenuation profiles of HA and CaOx at low energy thresholds of 15, 22, 29, and 36 keV and a varied tube potential of 50, 80 and 110 kVp were evaluated by a comparative contrast to noise ratio (CNR) to differentiate Type I and Type II μcals (Kirkbride *et al*. 2017). The attenuation of CaOx at a set concentration of 2000 mg/ml was compared to varying concentrations of HA (54.3, 211.7, 808.5, 1169.3 and 2000 mg/ml) within each energy bin. However, the attenuation profiles of HA and CaOx were difficult to differentiate because the overall attenuation of each is dominated by the calcium signal (Table 2). Therefore, the reliability of material separation was dependent on the concentration of the HA measured, with lower concentrations exhibiting less overlap in attenuation compared to CaOx. If applied to material decomposition, variability in the density of the μcal would change the accuracy of the material decomposition. The this study, we circumvented this problem by performing elemental decomposition of Ca, P, and O.

## 4. Conclusions

Utilizing multiple assumptions for calibration, photon-counting spectral CT and elemental decomposition by MLE in the image domain accurately identified and quantified spatially coincident compositions of Ca and P within three calcium crystal compositions (RMSE < 12%). Pathologically-relevant Type I (CaOx) and Type II (HA) μcals were accurately discriminated (AUC > 0.8) at a clinically-relevant size (max dimension < 1 mm). Thus, in spite of variation in material calibration, elemental decomposition in the image domain using photon-counting spectral CT enabled quantitative molecular imaging of calcium crystal compositions, for the discrimination of Type I and Type II μcals under conditions suitable for imaging, which has not been previously demonstrated. If translated to clinical spectral photon-counting mammography, the methods of this study have the potential to reduce the number of false-positives, unnecessary biopsy procedures, and overdiagnosis by mammographic screening..

## Acknowledgments

This research was supported by grants from the Kelly Cares Foundation, St. Joseph Health System, and the University of Notre Dame Equipment Renewal and Restoration Program. TEC was supported by the Martell Family Ph.D. Fellowship at University of Notre Dame. The authors acknowledge the Notre Dame Center for Environmental Science and Technology (CEST) for ICP-OES and the Notre Dame Integrated Imaging Facility (NDIIF).

